# Basal type I interferon signaling has only modest effects on neonatal and juvenile hematopoiesis

**DOI:** 10.1101/2022.07.18.500483

**Authors:** Yanan Li, Wei Yang, Helen C. Wang, Riddhi M. Patel, Emily B. Casey, Elisabeth Denby, Jeffrey A. Magee

**Affiliations:** Department of Pediatrics, Division of Hematology and Oncology, Washington University School of Medicine, 660 S. Euclid Ave, St. Louis, MO 63110; Department of Genetics, Washington University School of Medicine, 660 S. Euclid Ave, St. Louis, MO 63110

## Abstract

Type I interferon (IFN-1) regulates gene expression and hematopoiesis both during development and in response to inflammatory stress. We previously showed that during development, hematopoietic stem cells (HSCs) and multipotent progenitors (MPPs) induce IFN-1 target genes shortly before birth in mice. This coincides with the onset of a transition to adult hematopoiesis, and it drives expression of genes associated with antigen presentation. However, it is not clear whether perinatal IFN-1 modulates hematopoietic output, as has been observed in contexts of inflammation. We have characterized hematopoiesis at several different stages of blood formation, from HSCs to mature blood cells, and found that loss of the IFN-1 receptor (IFNAR1) leads to depletion of several phenotypic HSC and MPP subpopulations in neonatal and juvenile mice. Committed lymphoid and myeloid progenitor populations simultaneously expand. These changes had surprisingly little effect on production of more differentiated blood cells. Cellular Indexing of Transcriptomes and Epitopes by sequencing (CITE-seq) resolved the discrepancy between the extensive changes in progenitor numbers and modest changes in hematopoiesis, revealing stability in most MPP populations in *Ifnar1*-deficient neonates when the populations were identified based on gene expression rather than surface marker phenotype. Thus, basal IFN-1 signaling has only modest effects on hematopoiesis. Discordance between transcriptionally- and phenotypically-defined MPP populations may impact interpretations of how IFN-1 shapes hematopoiesis in other contexts, such as aging or inflammation.

**KEY POINTS:**

- Loss of type I Interferon signaling in neonatal mice depletes immature blood progenitors without compromising postnatal hematopoiesis
- Progenitor populations remain intact when measured by single cell transcriptomes rather than surface marker phenotypes

## INTRODUCTION

Inflammatory cytokines modulate hematopoiesis throughout life, particularly in response to infections or other inflammatory stressors.^1, 2^ Several cytokines, including interferon alpha (IFNα), interferon gamma (IFNγ) and interleukin-1beta (IL1β) all have been shown to drive quiescent hematopoietic stem cells (HSCs) into cycle and alter hematopoietic output.^3-6^ Acute exposure to type I interferons (IFN-1), which includes IFNα and interferon beta (IFNβ), promotes HSC proliferation and emergency megakaryopoiesis.^3, 7^ Likewise, IL1β exposure induces transient HSC proliferation and myelopoiesis.^5, 8^ These changes enhance blood production and immune function under conditions of stress. Inflammatory cytokines also help maintain basal hematopoietic output. For example, gut microflora stimulate IFNα and IL1β production, which in turn promotes myeloid progenitor expansion and myelopoiesis.^9, 10^ Over time, chronic cytokine stimulation promotes myeloid lineage bias that typifies aged HSCs.^9^ These observations show that infection and inflammation can shape hematopoietic output in adults by reprogramming immature progenitors.

Inflammatory cytokines also regulate fetal and neonatal hematopoiesis, though not necessarily in response to infectious stimuli. During mid-gestation in mice, IFNα, IFNγ and IL1β promote emergence of definitive HSCs from hemogenic endothelium in the dorsal aorta.^11-13^ During late gestation, IFN-1 levels spike and trigger expression of antiviral transcripts (e.g. *Ifit1, Oas2, Ifit3*), several transcription factors associated with IFN-1 signal transduction (*Stat1, Irf7* and *Irf9*), and major histocompatibility complex I (MHC-I) genes within HSCs and non-self-renewing multipotent progenitors (MPPs).^14^ The perinatal IFN-1 spike does not reflect a response to infection or a change in the fetal/maternal microbiome, as it occurs even in germ free mice.^14^ It coincides with the onset of a transition from fetal to adult transcriptional states in both HSCs and MPPs. Deleting the IFN-1 receptor, *Ifnar1*, slows the transition, indicating a causal interaction between sterile IFN-1 signaling and adult gene expression programs.^14^

It is less clear whether sterile IFN-1 signaling is necessary to support hematopoietic output during late gestation and in neonates. We previously showed that *Ifnar1* deletion prevents expansion of phenotypic MPP populations that takes place between embryonic day (E)16 and postnatal day (P)0.^14^ This finding aligns with recent observations in adult mice showing that *Stat1* deletion causes a reduction of MPP numbers.^10, 15^ However, *Ifnar1*-deficient mice develop normally, despite the aforementioned transcriptional changes, suggesting that the loss of phenotypic MPPs does not markedly impair hematopoiesis. Assessments of HSC and MPP frequencies by flow cytometry are confounded by the fact that Sca1, an HSC/MPP surface marker, is itself positively regulated by IFN-1.^3^ This raises the question of whether changes in phenotypic HSC, MPP and myeloid progenitor numbers in *Ifnar1*-or *Stat1*-deficient mice underlie functional changes that shape hematopoietic output, or whether they simply reflect changes in population surface marker phenotypes.

We have used *Ifnar1*-deficient mice to further characterize the role of IFN-1 signaling in neonatal and juvenile hematopoiesis. We found that *Ifnar1* deletion led to a pronounced decrease in phenotypic HSC and MPP numbers in newborn mice, with a concomitant increase in more mature myeloid and lymphoid progenitors. These changes were also evident in 14-day-old juvenile mice, and in adult mice after conditional *Ifnar1* deletion. Neonatal myelopoiesis, lymphopoiesis and blood production were unaffected by *Ifnar1* deletion despite the extensive changes within immature progenitor populations. Cellular Indexing of Transcriptomes and Epitopes by sequencing (CITE-seq)^16^ resolved this discrepancy. *Ifnar1* deletion had only minor effects on neonatal hematopoiesis, as defined by single cell transcriptomes, despite extensive changes in surface marker identity. We observed a modest decrease in HSC numbers and a modest decline in a T-lineage primed MPP population in newborn mice, but functional changes were minor relative to changes in phenotypic populations. The data indicate that non-infectious IFN-1 signaling induces transcriptional changes that promote antigen presentation and antiviral activity within HSCs/MPPs beginning in late gestation, but it has only modest effects on hematopoiesis.

## METHODS

### Mouse lines and husbandry

*Ifnar1-*null,^17^ *Ubc-CreER*,^*18*^ *and Ifnar1-*flox^19^ mice were obtained from the Jackson Laboratory. Mice were housed in a standard pathogen free barrier facility. For tamoxifen treatments, mice were fed 100 µg/gram body weight of tamoxifen in corn oil daily for 5 days by oral gavage. HSCs were cultured in Methocult and genotyped to confirm >90% *Ifnar1* deletion efficiency. All procedures were performed according to an IACUC approved protocol at Washington University School of Medicine.

### Flow cytometry

Cells were isolated, stained, analyzed and transplanted as previously described.^14, 20, 21^ Flow cytometry antibodies and surface marker phenotypes are defined in Table S1, and representative gating strategies are shown in Figures S2, S4 and S5. Flow cytometry was performed on a BD FACSAria Fusion flow cytometer (BD Biosciences).

### Cell cycle, BrdU Incorporation and Annexin V assays

For cell cycle assays, cells from the indicated populations were sorted directly into 70% methanol and stored at −20C for at least 24 hours. The cells were then washed in staining media, stained with propidium iodide/RNAse staining solution (Molecular Probes), and analyzed by flow cytometry. For BrdU incorporation assays, pregnant female mice were injected with BrdU (150 mg/kg body weight) on P19, 24-hours before cell collection. They were then placed on BrdU drinking water (1 mg/mL) and gave birth overnight. Livers were isolated from newborn pups for analysis. BrdU incorporation assays were performed as previously described using a BrdU flow kit (BD Pharminigen).^14^ For Annexin V assays, cells were stained for surface markers, as indicated above, as well as FITC or APC conjugated Annexin V dye (Biolegend).

### Western blot

LSK cells were isolated from P0 and 8-week-old adult mice, in parallel. Cells were then stimulated with IFNa (12.5 I.U./µL, Miltenyi) for 30 minutes, washed and then transferred to 10% trichloracetic acid. Western blots were then performed as previously described using antibodies from Cell Signaling Technologies (Table S1).^20^

### CITE-seq analysis

P0 liver cells from 3 pooled mice per genotype, or P14 bone marrow cells from 2-4 mice per genotype were c-kit selected and stained with tagged Total-Seq B antibodies (Biolegend) to CD150, CD48, CD135, CD34, CD201, CD117 (2B8 clone), CD16/32, CD127 and CD41. We isolated Lineage^-^c-Kit^+^ (LK) and Lineage^-^Sca1^+^c-Kit^+^ cells by flow cytometry and generated libraries with kits from 10x Genomics that included the Chromium Next GEM Single Cell 3′ kit v3.1 (PN-1000268), Chromium Single Cell 3’ Feature Barcode Library kit (PN-1000079), Dual Index kit NT Set A (PN-1000242) and Dual Index kit TT Set A (PN-1000215). All kits were used according to the manufacturer’s instructions. Resulting cDNA libraries were sequenced on an Illumina Novaseq S4.

The Cell Ranger v.6.0.1 pipeline (10x Genomics) were used to process data generated from the 10x Chromium platform. Digital gene expression (DGE) files were then filtered and normalized as previously described.^14^ We performed Iterative Clustering and Guide-gene Selection (ICGS, version 2) analysis with the AltAnalyze toolkit.^22^ Thresholds for surface barcode expression were set by visual inspection, and populations were defined based on phenotypes described in Table S2. Comparisons of population sizes were evaluated with a Chi-square test. For comparisons of gene expression between *Ifnar1*^*+/-*^ and *Ifnar1*^*-/-*^ progenitors, the expression data were stored as a Seurat data object and filtered to select cell that populated individual clusters based on the ICGS output. Differential expression analyses were then performed using the Seurat::FindMarkers function to compare across genotype groups with the Wilcoxon Rank Sum test.^23^

All CITE-seq data have been deposited into Gene Expression Omnibus as GSE205544.

### T cell potential assays

To measure T cell potential, OP9-delta cells were seeded in 96-well plates at 10,000 cells per well in MEMα, 20% fetal bovine serum and Penicillin-Streptomycin. Limiting does of Lineage^-^c-Kit^+^ cells (5, 20 or 35 cells) were sorted into at least 48 independent wells per genotype and cell dose. Cells were cultured with FLT3L and IL7 (12.5 ng/mL each for two weeks and then 2.5 ng/mL each for one week). Media was changed every 4 days. After 21 days, each well was analyzed for the presence of CD4 or CD8 T cells by flow cytometry. Extreme Limiting Dilution Analysis (ELDA) was used to calculate the fraction of LK cells with T cell potential, and to compare *Ifnar1*^*+/-*^ and *Ifnar1*^*-/-*^ mice.

## RESULTS

### *Ifnar1* deletion alters phenotypic HSC, MPP and committed progenitor populations in neonatal and juvenile mice without disrupting hematopoiesis

We used *Ifnar1*-deficient mice to thoroughly characterize the effects of IFN-1 signaling on neonatal and juvenile hematopoiesis. Neonatal and adult HSCs/MPPs are similarly responsive to IFN-1, as measured by STAT1 and STAT3 phosphorylation (Figure S1), suggesting that IFN-1 signal transduction does not change with age, but the late gestational IFN-1 pulse could nevertheless alter postnatal hematopoiesis. We used nomenclature laid out by Pietras et al. to measure phenotypic HSC and MPP subpopulations in neonatal liver (postnatal day (P)0) and juvenile bone marrow (P14) (Figure S2A, Table S2). These included long-term HSCs (LT-HSCs) that have extensive self-renewal capacity, short-term HSCs (ST-HSCs) that have limited self-renewal capacity, MPP2s that demonstrate erythroid and megakaryocyte bias, MPP3s that demonstrate myeloid bias and MPP4s that demonstrate lymphoid bias. To simplify breeding strategies, *Ifnar1*^*+/-*^ were used as controls rather than *Ifnar1*^*+/+*^ mice. We confirmed that *Ifnar1*^*+/+*^ and *Ifnar1*^*-/-*^ mice had equivalent numbers of HSCs, MPPs and myeloid progenitors (Figure S3).

We previously showed that *Ifnar1*^*-/-*^ mice have normal HSC and MPP numbers at E16, but that MPP numbers are reduced in *Ifnar1*^*-/-*^ mice by P0.^14^ We reaffirmed the previous findings, observing depletion of several phenotypic HSC and MPP subpopulations in livers of P0 *Ifnar1*^*-/-*^ mice (Figure 1A-E). Significant, but less severe, reductions in HSC/MPP numbers were observed in P14 juvenile mice (Figure 1F-J). We also conditionally deleted *Ifnar1* in adulthood by administering tamoxifen to *Ubc-CreER; Ifnar1*^*f/f*^ mice at 6-weeks-old and then evaluating HSC and MPP numbers 4 weeks later. All MPP subpopulations (MPP2, MPP3 and MPP4) were significantly reduced in *Ifnar1*-deleted mice, whereas HSC numbers were normal (Figure 1K-O). Altogether, the data show that *Ifnar1* deletion reduces phenotypic MPP numbers across a range of developmental stages.

**Figure 1.**
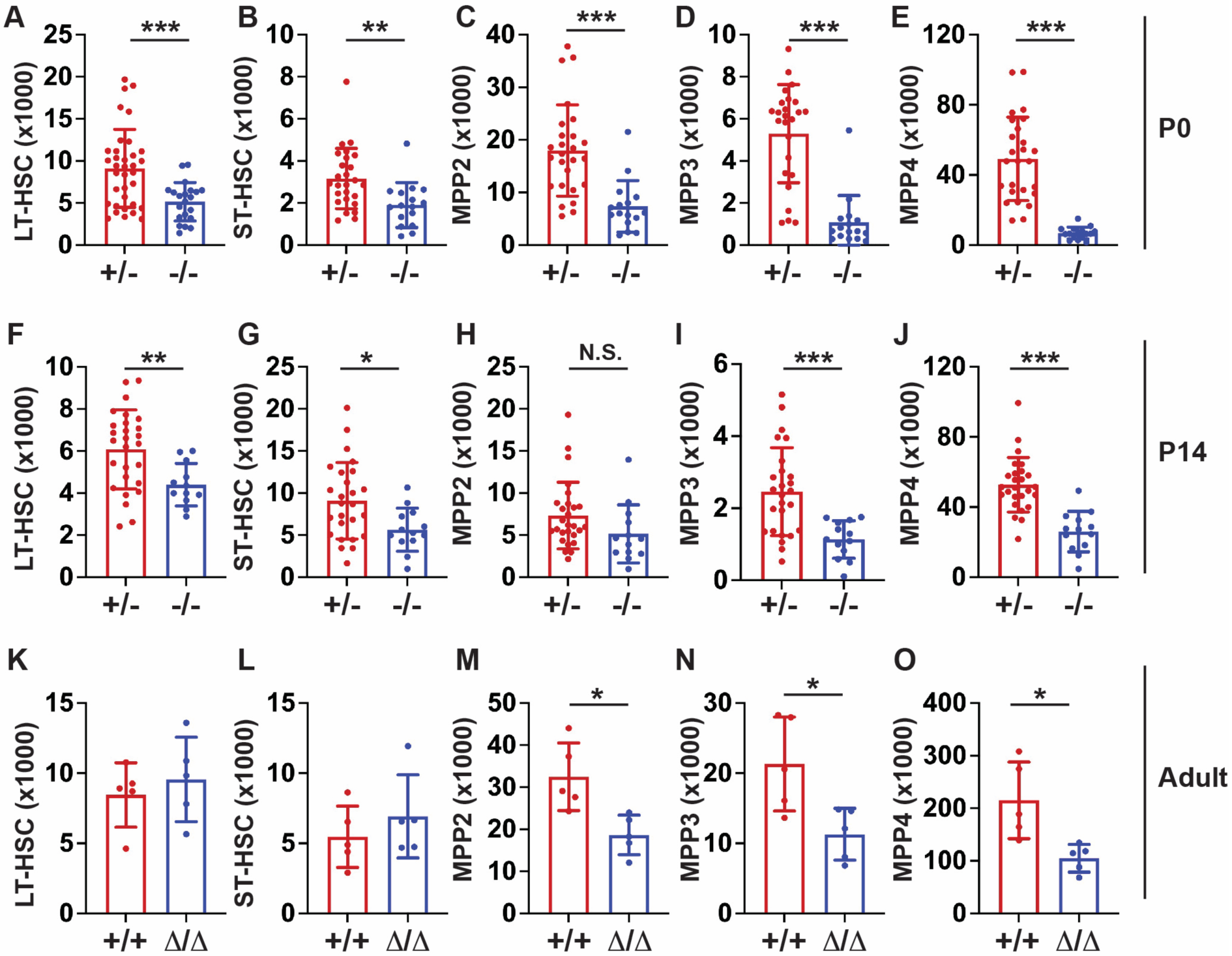
*Ifnar1* deletion leads to reductions in phenotypic HSCs and MPPs during neonatal, juvenile and adult stages of life. (A-E) Numbers of indicated HSC and MPP subpopulations in livers of P0 *Ifnar1*^*+/-*^ and *Ifnar1*^*-/-*^ mice, n=21-35. (F-J) Numbers of indicated HSC and MPP subpopulations in bone marrow (two hindlimbs) of P14 *Ifnar1*^*+/-*^ and *Ifnar1*^*-/-*^ mice, n=12-27. (K-O) Numbers of indicated HSC and MPP subpopulations in bone marrow of 10-week-old control (Cre-negative) and Ubc-CreER; *Ifnar1*^*f/f*^ mice after tamoxifen treatment at 6 weeks old, n=5. For all panels, error bars reflect standard deviation. Surface marker phenotypes are specified in Table S2 and gating strategies are shown in Figure S2. *p<0.05, **p<0.01, ***p<0.001 by two-tailed Student’s t-test.

In light of the marked depletion of HSCs and MPPs in *Ifnar1*-deficient mice, we next evaluated whether more mature stages of blood production are similarly disrupted. We measured pre-granulocyte-monocyte progenitor (pGM), granulocyte-monocyte progenitor (GMP), pre-megakaryocyte progenitor (pre-Meg), megakaryocyte progenitor (MkP), common lymphoid progenitor (CLP) and pre-erythroid colony forming unit (pre-CFU-E) numbers at neonatal (P0) and juvenile (P14) stages using surface marker phenotypes described by Pronk et al. (Figure S2B and C; Table S2). We also measured more mature blood cells, including B- and T-cell progenitors and fully differentiated peripheral blood cells. We observed significant expansion, rather than depletion, of pGM, GMP, pre-Meg, MkP and CLP populations (Figure 2), particularly at P0 when IFN-1 target gene expression is highest. These increases in population size were paradoxical, given the depletion of HSC and MPP precursors. Interestingly, *Ifnar1* deletion neither impaired or enhanced production of mature blood progenitors or fully differentiated blood cells, as B-cell progenitors (Figure 3A-D), T-cell progenitors (Figure 3E-H) and peripheral blood counts were all normal in *Ifnar1*^*-/-*^ mice at P0 and P14 (Figure 3I-L). Thus, profound shifts in the composition of immature HSC and MPP populations in *Ifnar1*^*-/-*^ neonates does not result in altered hematopoietic output.

**Figure 2.**
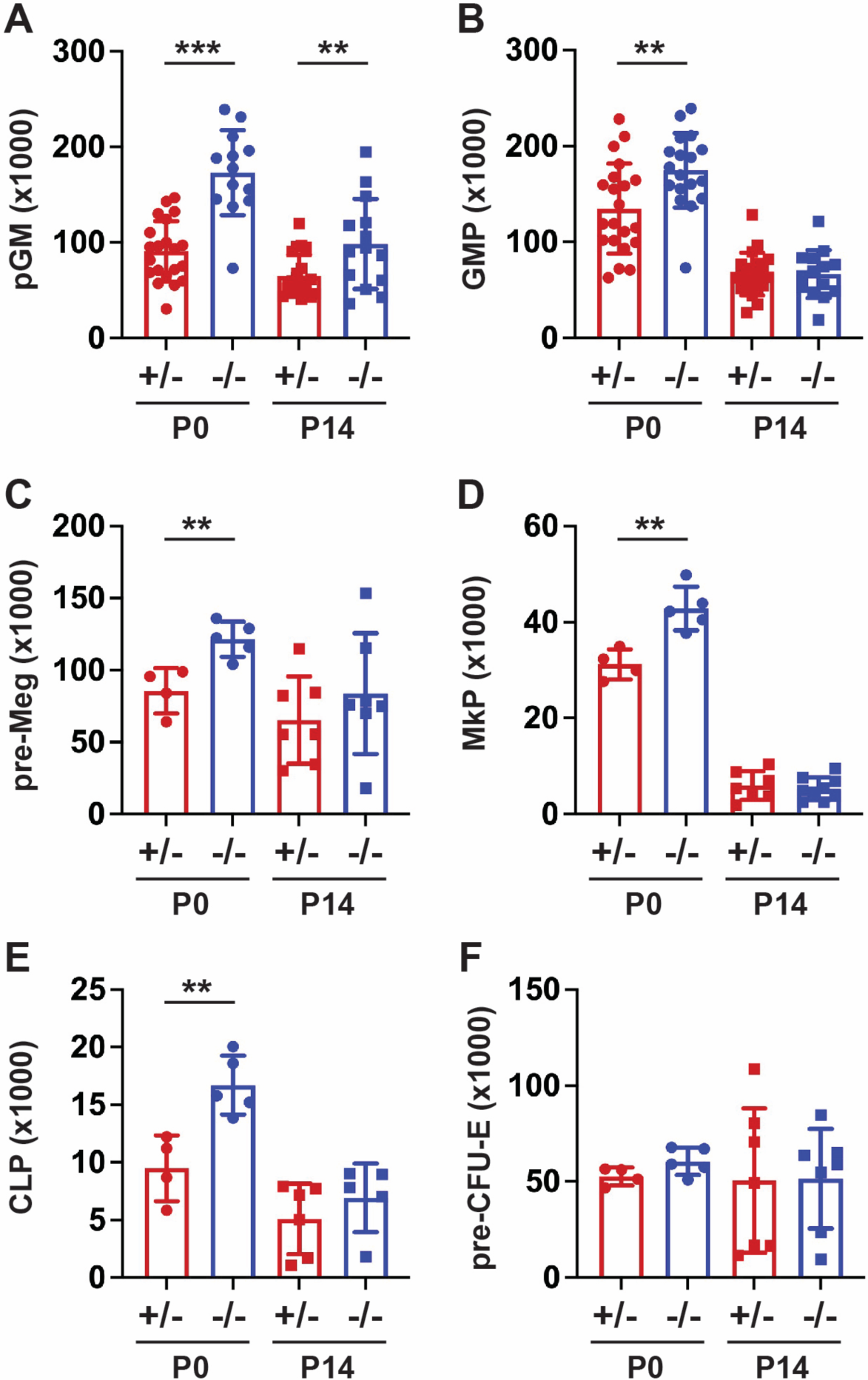
*Ifnar1* deletion leads to expansion of phenotypic committed myeloid and lymphoid progenitor populations in neonates and juveniles. Absolute numbers of committed myeloid (A, B), megakaryocyte (C, D), lymphoid (E) and erythroid (F) progenitors in P0 liver or P14 bone marrow from mice of *Ifnar1*^*+/-*^ and *Ifnar1*^*-/-*^ mice. For panels A and B, n=13-21. For panels C-F, n=4-7. For all panels, error bars reflect standard deviation. Surface marker phenotypes are specified in Table S2 and gating strategies are shown in Figure S2. **p<0.01, ***p<0.001 by two-tailed Student’s t-test.

**Figure 3.**
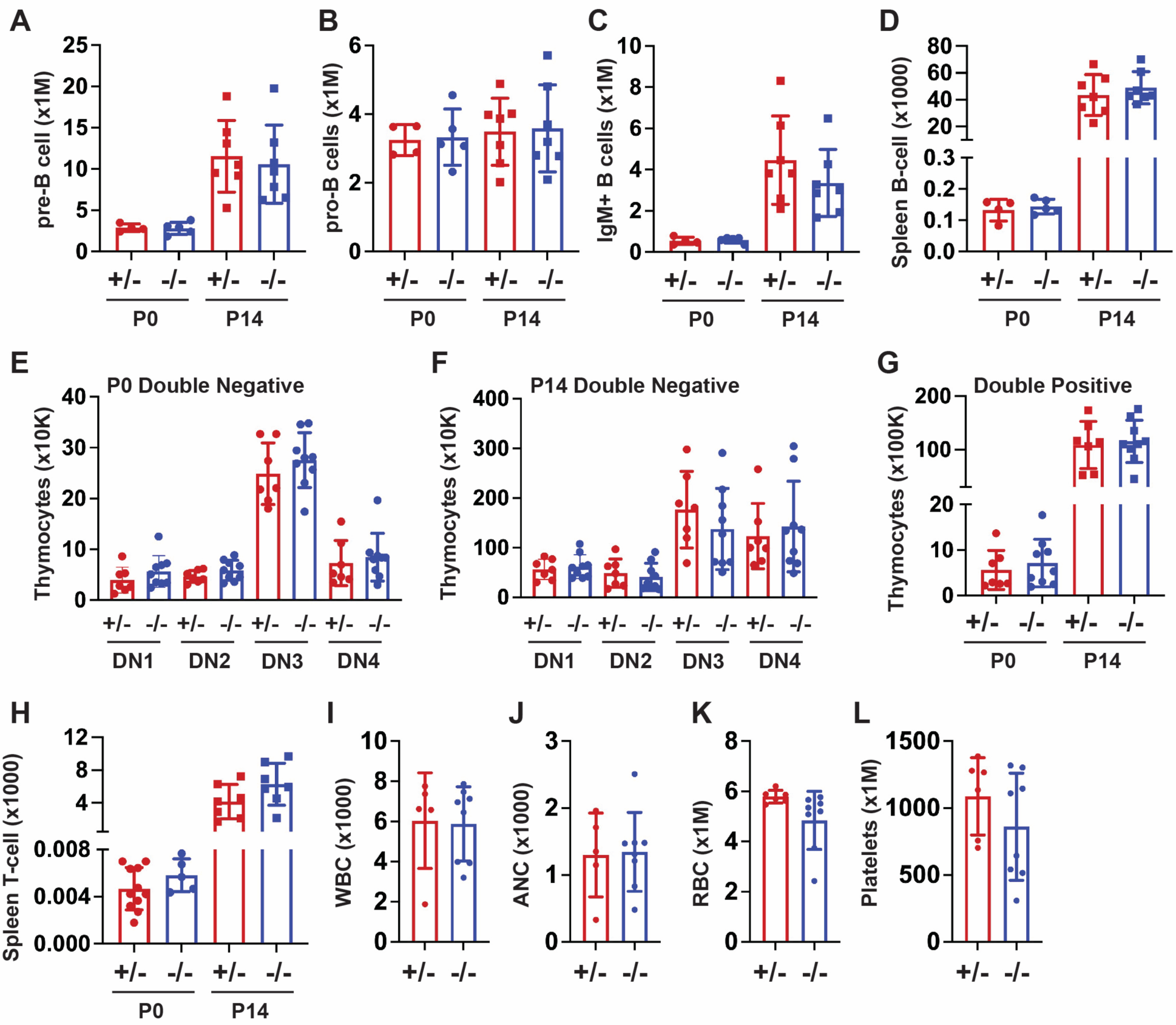
*Ifnar1* deletion does not impair lymphopoiesis or mature blood production. (A-D) Absolute numbers of indicated B cell progenitor populations in P0 liver or P14 bone marrow from mice of *Ifnar1*^*+/-*^ and *Ifnar1*^*-/-*^ mice, or spleen B cell populations from the same mice. (E-H) Absolute numbers of thymocyte subpopulations *Ifnar1*^*+/-*^ and *Ifnar1*^*-/-*^ mice, or spleen T cell populations from the same mice. (I-L) White blood cell (H), absolute neutrophil (I), red blood cell (J) and platelet (K) counts presented for 1 µL of blood from P14 mice. For all panels, n=4-11 and error bars reflect standard deviation. Surface marker phenotypes are specified in Table S2 and gating strategies are shown in Figure S4. Comparisons performed by two-tailed Student’s t-test did not show significant differences between *Ifnar1*^*+/-*^ and *Ifnar1*^*-/-*^ mice for any measurement.

### Basal IFNγ and IL1β signaling do not modulate neonatal progenitor population sizes

In addition to IFN-1, IL1β and IFNγ have both been shown to regulate fetal HSC emergence and adult HSC/MPP stress responses.^11-13^ We therefore tested whether either cytokine could modulate HSC/MPP numbers, alone or in cooperation with IFN-1. We generated mice deficient for either the IL1β receptor (*Il1r1*) or the IFNγ receptor (*Ifngr1*). In each case, the loss-of-function alleles were crossed with *Ifnar1*-deficient mice to test for genetic interactions between the inflammatory cytokine receptors. *Il1r1* deletion did not lead to HSC or MPP depletion in P0 mice, nor did it amplify effects of *Ifnar1* deletion (Figure 4A-E). Likewise, *Ifngr1* deletion did not lead to HSC or MPP depletion in P0 mice by itself, nor did it amplify effects of *Ifnar1* deletion (Figure 4F-J). These findings show that IFN-1 is selectively responsible for expansion of phenotypic HSC and MPP populations in neonates.

**Figure 4.**
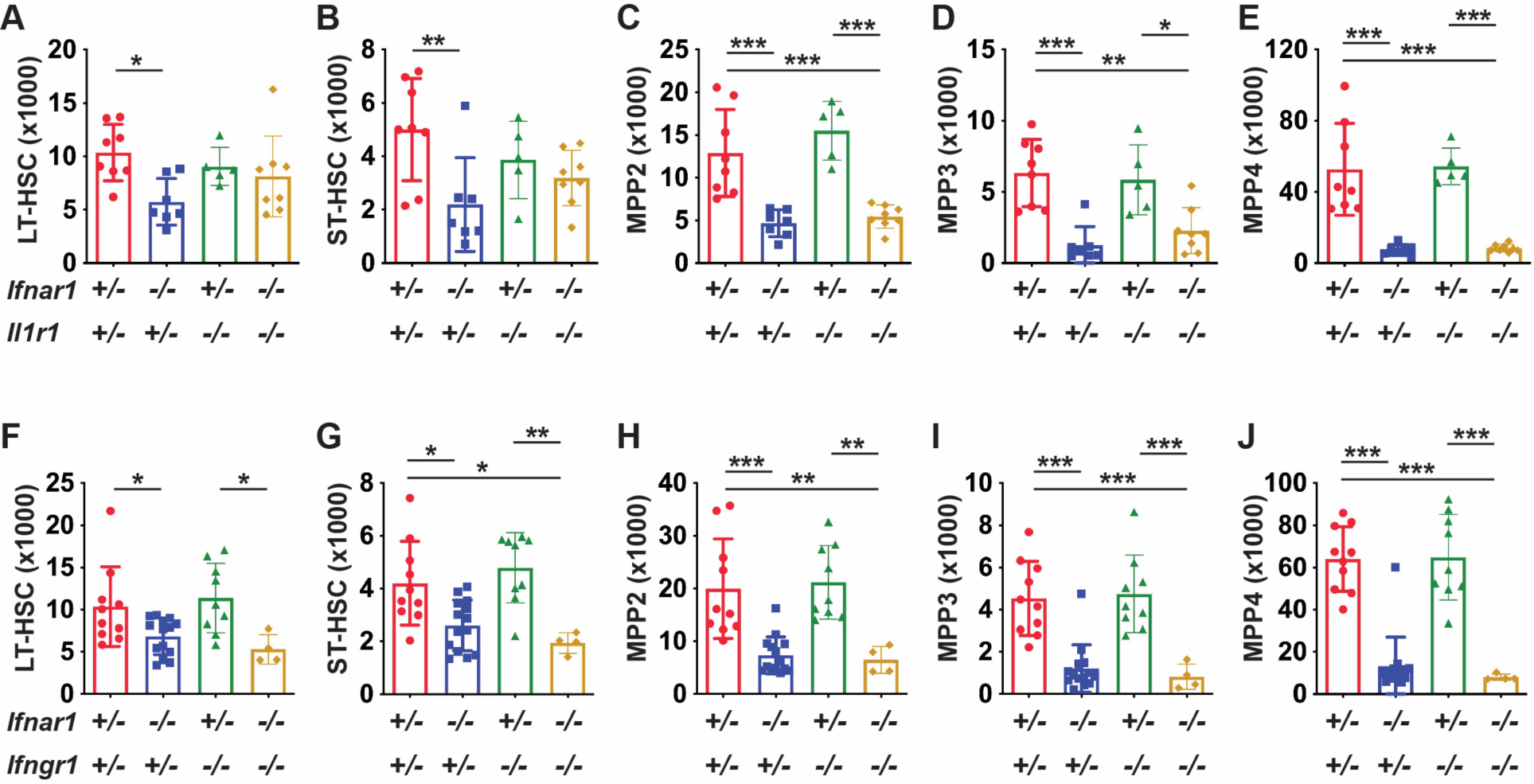
IL-1 and IFNγ signaling is not required to sustain neonatal HSC/MPP populations. (A-E) Numbers of indicated HSC and MPP subpopulations in livers of P0 mice with heterozygous or homozygous deletions of *Il1r* and *Ifnar1*, n=4-14. (F-J) Numbers of indicated HSC and MPP subpopulations in livers of P0 mice with heterozygous or homozygous deletions of *Ifngr1* and *Ifnar1*, n=4-14. For all panels, error bars reflect standard deviation. *p<0.05, **p<0.01, ***p<0.001 by one-way ANOVA with Holm-Sidak posthoc test to correct for multiple comparisons.

### IFN-1-dependent changes in neonatal MPP population sizes do not reflect changes in proliferation or cell death

We sought to understand why HSC and MPP populations decline in *Ifnar1*^*-/-*^ neonates, even as committed progenitor populations expand. We tested whether these changes reflect differences in cell proliferation rates or cell death. To measure proliferation, we sorted HSCs (LT- and ST-HSCs), MPPs (MPP2, MPP3 and MPP4), pGMs and GMPs from P0 livers into methanol. We then performed propidium iodide staining to assess the fraction of each population in S/G2/M phase of the cell cycle (Figure S5A). *Ifnar1*^*-/-*^ MPPs showed a modest but significant increase, rather than the anticipated decrease, in proliferation relative to controls (Figure 5B). The remaining populations showed no differences in proliferation between *Ifnar1*^*+/-*^ and *Ifnar1*^*-/-*^ mice (Figure 5A, C and D). We confirmed that P0 *Ifnar1*^*-/-*^ MPPs proliferate at a slightly higher rate than *Ifnar1*^*+/-*^ controls based on 5’-bromo-2’-deoxyuridine (BrdU) incorporation (Figure 5E, Figure S5B). However, we did not observe a similar increase in proliferation, as measured by propidium iodide, in P14 MPPs (Figure 4F). There were no *Ifnar1*-dependent differences in programmed cell death rates in P0 HSCs, MPPs, pGMs or GMPs as measured by Annexin V staining (Figure 5G, Figure S5C, D). Altogether, the data indicate that changes in proliferation and cell death do not adequately account for the reduction in HSCs/MPPs and the increase in pGMs/GMPs observed in newborn *Ifnar1*^*-/-*^ mice.

**Figure 5.**
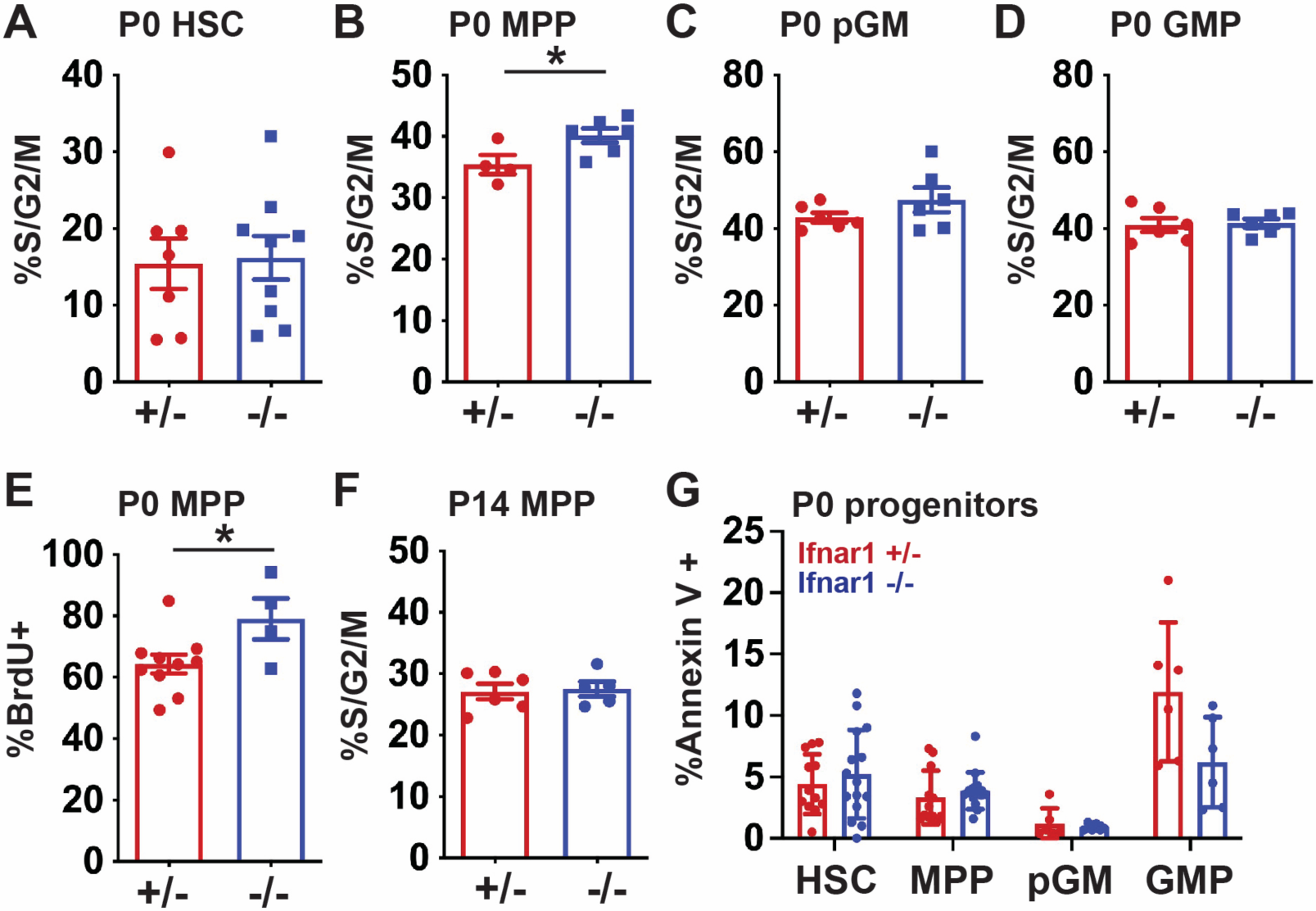
*Ifnar1* deletion does not impair proliferation or promote death or neonatal HSCs or MPPs. (A-D) Percentages of P0 HSCs (CD150^+^CD48^-^LSK), MPPs (CD48^+^LSK), pGMs or GMPs in S/G2/M phase of the cell cycle in *Ifnar1*^*+/-*^ and *Ifnar1*^*-/-*^ mice, as measured by propidium iodine staining, n=4-9. (E) Percent of BrdU positive MPPs (CD48^+^LSK) after a 24-hour pulse between E19 and P0, n=4-10. (F) Percent of P14 MPPs (CD48^+^LSK), in S/G2/M phase of the cell cycle as measured by propidium iodine staining n=5-6. (G) Percent of Annexin V+, DAPI-negative apoptotic HSCs, MPPs, pGM and GMP in the P0 livers of *Ifnar1*^*+/-*^ and *Ifnar1*^*-/-*^ mice, n=6-15. For all panels, error bars reflect standard deviation. Representative gating strategies are shown in Figure S5. *p<0.05 by two-tailed Student’s t-test.

### *Ifnar1* deletion has only modest effects on progenitor populations as defined by single cell transcriptomes, in contrast to extensive changes observed by surface phenotype

We used CITE-seq to better characterize effects of IFN-1 signaling on neonatal and juvenile hematopoiesis, independent from surface marker phenotypes. The goal was to understand why committed progenitor populations paradoxically expand in *Ifnar1*^*-/-*^ mice while HSC and MPP populations decline, and why changes in immature progenitor populations do not alter hematopoietic output. The CITE-seq assays incorporated feature barcodes for nine different surface markers, thus enabling us to resolve HSC, MPP, pGM, GMP and CLP subpopulations while simultaneously assessing single cell gene expression (Figure S6). We obtained transcriptomes for Lineage^-^c-Kit^+^ (LK) and Lineage^-^Sca1^+^c-Kit^+^ (LSK) cells and generated libraries independently. We then aggregated the data after sequencing. We took this approach so that we could compare changes in progenitor population frequencies when Sca1 was both included and not included as a selection marker, given that Sca1 is an IFN-1 target gene. We used Iterative Clustering with Guide-gene Selection, version 2 (ICGS2)^22, 24^ to cluster the data and overlay surface marker phenotypes (Figure 6A, B; Tables S2 and S3). The CITE-seq data redemonstrated depletion of HSC and MPP populations, and concomitant expansion of CLP, pGM and GMP populations, in *Ifnar1*^*-/-*^ mice (Figure 6B).

**Figure 6.**
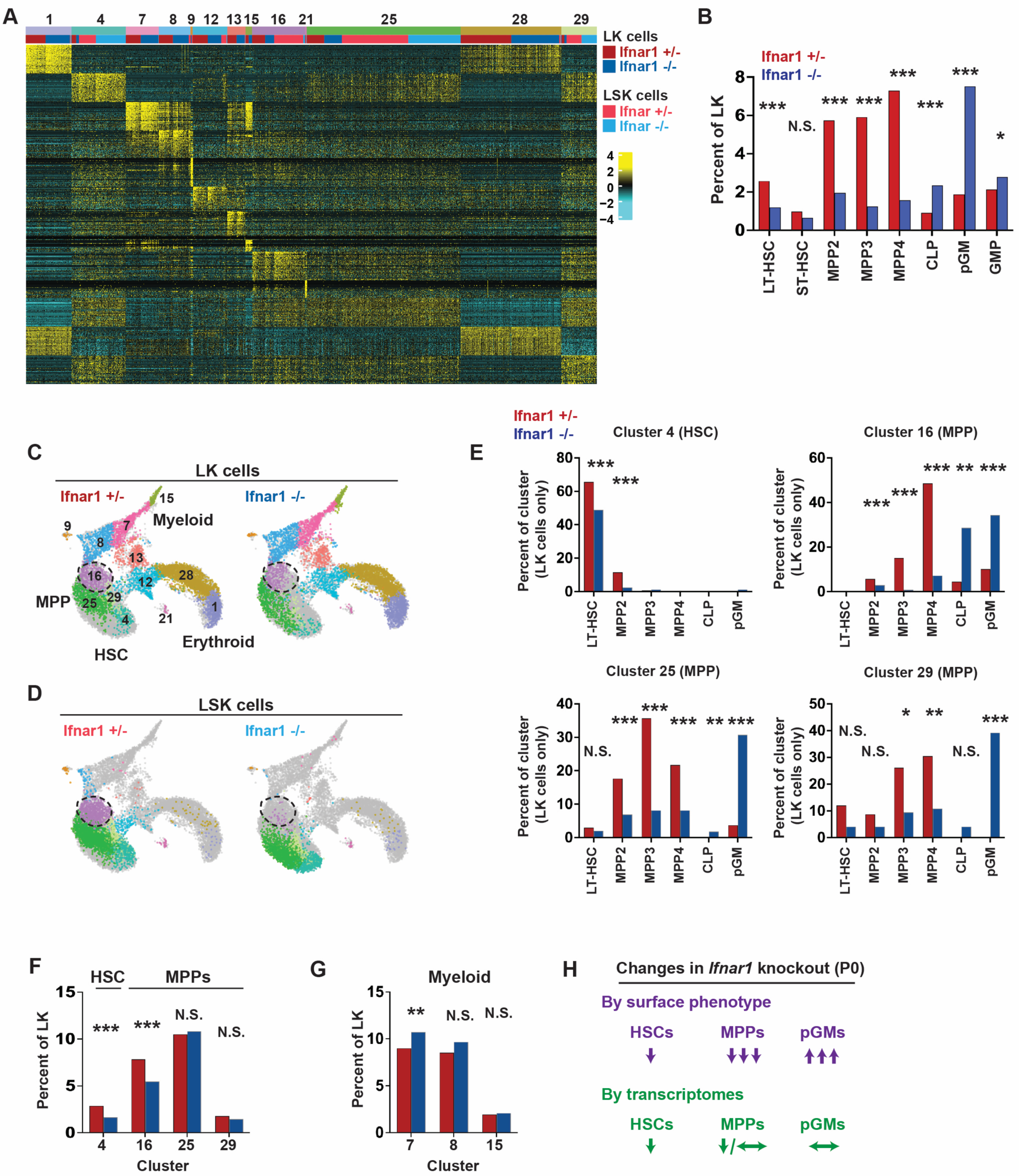
*Ifnar1* deletion has only modest effects on immature progenitor populations when the populations are defined by single cell transcriptomes rather than surface marker phenotypes. (A) Heatmap identifying 13 distinct clusters after iterative clustering via ICGS2. The clusters are numbered (ranging from 1-29) as defined by the ICGS2 algorithm. The clustering reflects aggregates of separately sorted LK and LSK cells from *Ifnar1*^*+/-*^ and *Ifnar1*^*-/-*^ mice. Genotypes and cell types are indicated by color coding in the second horizontal bar above the heatmap. (B) HSC, MPP and committed progenitor populations were defined based on surface markers as per Table S2. The frequencies of each phenotypically-defined population within the LK samples are shown for *Ifnar1*^*+/-*^ and *Ifnar1*^*-/-*^ mice. (C, D) UMAP plots showing clustering results for *Ifnar1*^*+/-*^ and *Ifnar1*^*-/-*^ LK and LSK cells. Differentiation states are indicated based on marker gene expression for each cluster. Clusters are color coded to match panel A. Cluster 16, a T cell biased MPP population, is indicated with the hashed circle. (E) Percentages of LK cells in clusters 4, 16, 25 and 29 with the indicated surface marker phenotypes. (F) Percentages of LK cells in HSC cluster 4 or MPP clusters 16, 25 and 29 in *Ifnar1*^*+/-*^ and *Ifnar1*^*-/-*^ mice. (G) Percentages of LK cells in myeloid clusters 7, 8 and 15 in *Ifnar1*^*+/-*^ and *Ifnar1*^*-/-*^ mice. (H) Summary of changes in HSC, MPP and pGM frequencies when the populations are defined by surface marker phenotype as compared to single cell transcriptomes. *p<0.05, **p<0.01, ***p<0.001 by the Chi-square test.

We next tested whether frequencies of transcriptionally-defined progenitor populations shifted in *Ifnar1*^*-/-*^ neonates. ICGS2 identified 13 unique clusters (Figure 6A, C, D). Cluster 4 expressed guide genes consistent with LT-HSC identity (Table S4, Figure 7A), and a majority of cells within the cluster bore LT-HSC surface markers (CD201^+^CD150^+^CD48^-^LSK) in wildtype mice (Figure 6E). The phenotypic LT-HSC population declined within cluster 4 in *Ifnar1*^*-/-*^ mice. Note that for this analysis, we focused exclusively on the LK cell data so that cells that downregulate Sca1 in *Ifnar1*-deficient mice would still be included in the analysis. We also identified three clusters – numbers 16, 25 and 29 – with guide gene profiles and surface marker phenotypes indicative of MPPs (Figure 6C-E). These clusters, along with cluster 4, accounted for almost all LSK cells (Figure 6D), consistent with MPP identities. Cells within clusters 16, 25 and 29 shifted from MPP to CLP and pGM identities in *Ifnar1*^*-/-*^ mice (Figure 6E). IFN-1 target gene expression was reduced in clusters 16, 25 and 29 consistent with our prior studies (Table S5), but only cluster 16 was modestly reduced in size (Figure 6F). Myeloid progenitor clusters 7, 8 and 15 showed little to no expansion despite increases in phenotypic pGMs (Figure 6G). A similar analysis at P14 revealed only modest changes in MPP population frequencies, as defined by gene expression (Figure S7A-D). Thus, IFN-1 is not necessary to sustain neonatal or juvenile MPPs when one uses gene expression rather than surface markers to identify the populations (Figure 6H). The simplest explanation for this phenomenon is that Sca1 expression declines in *Ifnar1*-deficient MPPs and causes them to appear as CLPs and pGMs.

**Figure 7.**
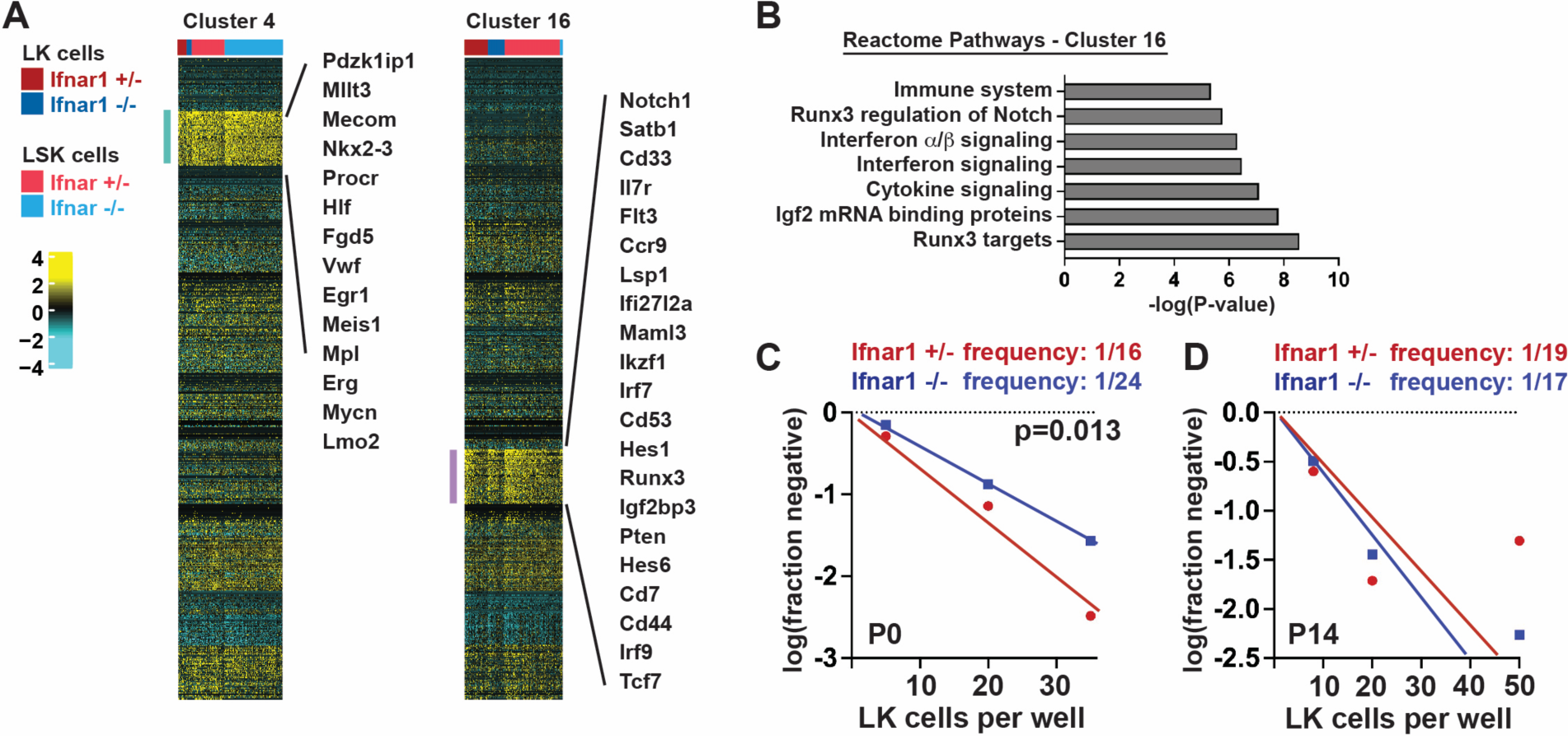
*Ifnar1* deletion causes modest reduction in T cell potential in neonatal mice. (A) Clusters 4 and 16 were modestly depleted in *Ifnar1*^*-/-*^ neonates as compared to *Ifnar1*^*+/-*^ neonates. Cluster 4 cells expressed genes consistent with HSC identity. Cluster 16 cells expressed some genes associated with myeloid bias (e.g. *Cd33*) and many genes associated with T lymphoid bias (e.g. Notch, Hes1, Runx3, etc.). (B) Reactome pathway analysis of marker genes for cluster 16 showed interferon-related genes as well as Notch/Runx3-associated genes that would align with T cell bias. (C, D) Results from limit dilution assays of LK cells plated on OP9-delta stroma to assess T cell potential at P0 and P14. The frequencies of LK cells with T cell potential in *Ifnar1*^*+/-*^ and *Ifnar1*^*-/-*^ mice were calculated and compared with ELDA, n=48-96 wells per cell dose and genotype.

### IFN-1 sustains a T cell-biased MPP population in neonates

As noted above, we observed modest decreases in HSCs (cluster 4) and one MPP population (cluster 16) in *Ifnar1*^*-/-*^ neonates as a fraction of the LK population (Figure 6F, Figure 7A). We have previously shown that *Ifnar1*-deficient HSCs function normally in repopulating assays and therefore did not characterize them further. Instead, we sought to characterize the function of MPPs that populated cluster 16. Guide genes for cluster 16 included *Notch1, Il7r, Runx3, Pten* and several Notch signal transduction intermediates (*Maml3, Hes1, Hes6*), all suggesting a T lineage bias (Figure 7A, B). To test whether T-cell potential was impaired in *Ifnar1*^*-/-*^ neonates, we plated limiting numbers of LK cells on OP9-delta stromal cells and counted wells that gave rise to T cells. We observed a 50% reduction in T-cell potential in P0 *Ifnar1*^*-/-*^ LK cells, consistent with the changes observed by CITE-seq (Figure 7C). We identified a population with similar gene expression in P14 mice (cluster 10; Figure S7; Table S4). This cluster was not significantly depleted in P14 *Ifnar1*^*-/-*^ LK cells (Figure S7B), and T cell potential was not impaired at P14 (Figure 7). Thus, IFN-1 signaling helps sustain a subset of T cell-biased MPPs specifically in neonates.

## DISCUSSION

Inflammatory cytokines have well-established roles in promoting myelopoiesis and megakaryopoiesis under stress conditions,^1, 2^ but their roles in baseline hematopoiesis are just starting to emerge. In this paper, we have carefully evaluated the effects of basal IFN-1 signaling on neonatal and juvenile hematopoiesis. Consistent with prior work,^14^ deleting the IFN-1 receptor (*Ifnar1*) had profound effects on many immature progenitor populations, at least based on surface marker phenotypes. However, despite extensive efforts, we found that *Ifnar1* deletion does not impair mature blood cell production in neonates or adults. Furthermore, depletion of MPPs in *Ifnar1*-deficient neonates appears to reflect shifts in surface marker expression rather than loss of the populations altogether, given that transcriptionally-defined MPP populations remained largely intact in our CITE-seq assays (Figure 6). We did observe a modest decline in one T cell-biased MPP population at P0 that correlated with a reduction in T cell potential (Figure 7). Transcriptionally defined HSCs also declined in *Ifnar1*-deficient neonates, but our prior studies showed that HSC function remains intact.^14^ Altogether, the data do not support a prominent role for physiologic IFN-1 signaling in sustaining neonatal or juvenile hematopoiesis, and other cytokines (e.g., IFNγ and IL1β) also appeared to have negligible effects in the absence of infection (Figure 4).

Our findings pertain specifically to IFN-1 signaling during normal ontogeny, and they do not address conditions of stress or aging. Maternal infections could alter postnatal blood production, and our data are silent on that context. Likewise, hematopoietic changes in response to microbial colonization and infection in adulthood may be different than responses to sterile IFN-1 signaling during fetal/neonatal stages. Nevertheless, our CITE-seq data indicate that even under basal conditions, IFN-1 signaling can have profound effects on HSC/MPP surface marker phenotypes. Changes in Sca1 expression that have previously been shown to accompany hyper-inflammation^3^ can also impact HSC/MPP phenotypes in non-inflamed states. Thus, efforts to use surface markers to characterize hematopoiesis under germ-free or cytokine-deficient contexts should be made with some caution. CITE-seq and other single cell techniques can provide valuable complementary insights.

Our findings raise an important question: What is the role of the perinatal IFN-1 spike if it does not modulate hematopoietic output? One key function is to promote expression of genes associated with antigen presentation and antiviral responses, particularly Major Histocompatibility Complex (MHC) I genes, within the postnatal HSC and MPP populations. We described these changes previously and redemonstrate them here.^14^ Interferon target gene expression has also been shown to increase between mid-gestation in human HSCs/MPPs, suggesting that the late-gestation IFN-1 response may be conserved.^25, 26^ Our data suggest that perinatal IFN-1 may prime postnatal progenitors to engage the adaptive immune system and activate antiviral programs without substantially influencing hematopoietic output. It is also possible that flux through the HSC/MPP/pGM populations changes in response to IFN-1 signaling, even if the populations themselves remain stable in size. For example, it has recently been proposed that MPPs, rather than HSCs, sustain most postnatal blood production into early adulthood.^27, 28^ The ratio of HSC- and MPP-dependent blood production could shift in response to IFN-1 or other inflammatory cytokines. Future lineage tracing studies can resolve this question.

## ACKNOWLEDGEMENTS

This work was supported by grants to J.A.M. from the NHLBI (R01 HL152180 and R01 HL136504), Alex’s Lemonade Stand Foundation (‘A’ Award), Hyundai Hope on Wheels, the Alvin J. Siteman Cancer Center Investment Program (supported by the Foundation for Barnes-Jewish Hospital and NCI Cancer Center Support Grant P30 CA091842), and the Children’s Discovery Institute of Washington University and St. Louis Children’s Hospital. J.A.M. is a Leukemia and Lymphoma Society Scholar.

## AUTHOR CONTRIBUTIONS

J.A.M. designed and oversaw all experiments, conducted experiments, interpreted data, wrote the manuscript, and secured funding. Y.L. conducted experiments, interpreted data and wrote the manuscript. W.Y. performed all bioinformatic analyses. H.C.W., R.M.P., E.B.C. and E.D. performed experiments and interpreted data. All authors reviewed and edited the manuscript. Correspondence and requests for materials should be addressed to.

## DISCLOSURE OF CONFLICTS OF INTEREST

The authors declare no competing interests.

## Notes

### Competing Interest Statement

The authors have declared no competing interest.

