## Supplemental Figures and legends for "Basal type I interferon signaling has only modest effects on neonatal and juvenile hematopoiesis"

#### SUPPLEMENTARY TABLES

**Table S1. Antibodies used for flow cytometry, Western blots and CITE-seq.** The CITE-seq indices were used to align single cell transcriptomes with surface marker expression.

**Table S2. Surface marker phenotypes used to define progenitor populations in flow cytometry and CITE-seq assays.**

**Table S3. Numbers of cells passing quality control, genes per cell and unique molecular indices per cell for CITE-seq experiments.**

**Table S4. Cluster-specific marker genes as defined by ICGS2.**

**Table S5. Differentially expressed genes from control and Ifnar1-deficient HSC and MPP clusters at P0.** A negative log2 fold change indicates genes that show reduced expression in *Ifnar1*$^{-/-}$ cells relative to *Ifnar1*$^{+/-}$ cells from the same cluster. Ribosomal and mitochondrial genes were regressed from the analysis.

#### Figure S1

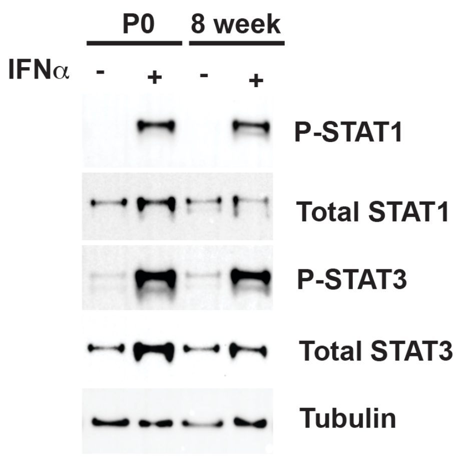


**Figure S1. Neonatal and adult HSCs/MPPs respond similarly to IFN-1 stimulation.** The Western blot shows STAT1 and STAT3 phosphorylation in P0 or 8-week-old LSK cells after exogenous stimulation with IFNα (12.5 units/μL for 30 minutes).

### Figure S2

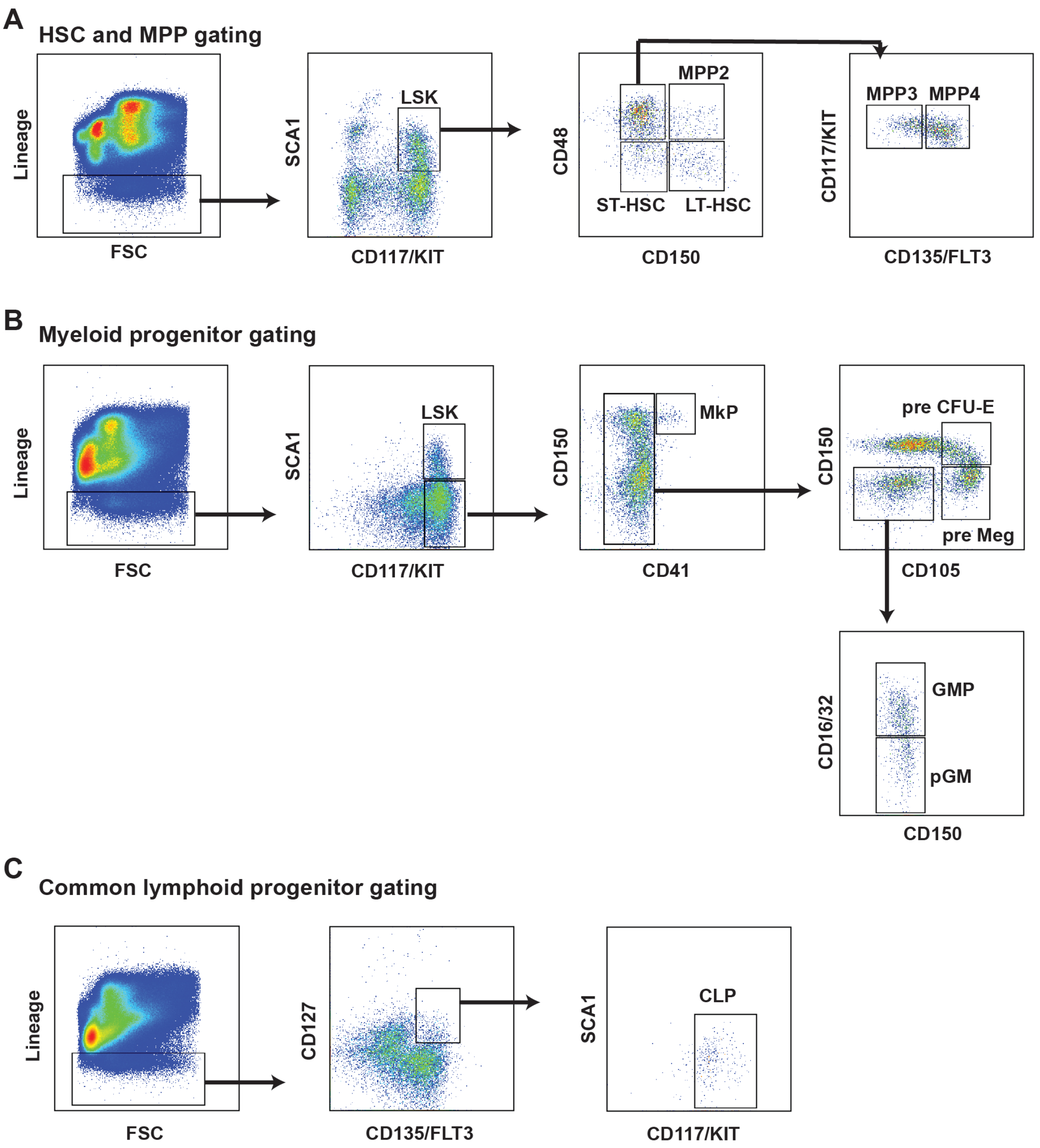


**Figure S2. Gating strategies for HSC, MPP and committed progenitor populations.** (A) Representative gating for neonatal for HSC and MPP subpopulations. (B) Representative gating for neonatal myeloid, megakaryocytic and erythroid progenitor populations. (C) Representative gating for neonatal for common lymphoid progenitors.

#### Figure S3

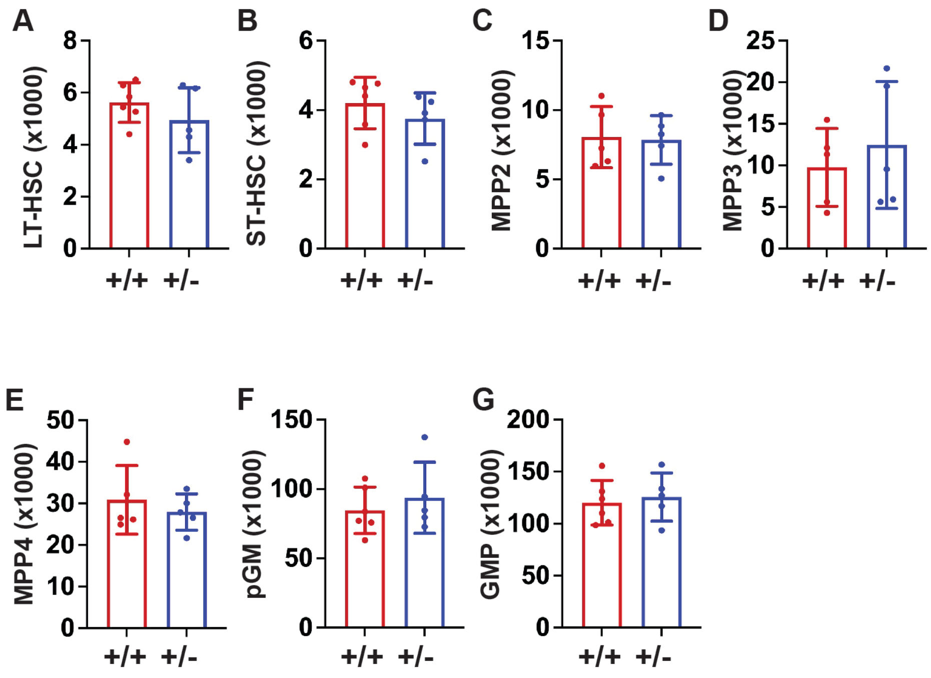


**Figure S3. Progenitor numbers in *Ifnar1*$^{+/+}$ and *Ifnar1*$^{+/-}$ neonates are indistinguishable.** Numbers of the indicated cell populations are shown in P0 livers for mice of the indicated genotypes. For all panels, n=5-6 and error bars reflect standard deviation. Comparisons performed by two-tailed Student's t-test did not show significant differences between *Ifnar1*$^{+/+}$ and *Ifnar1*$^{+/-}$ mice for any measurement.

#### Figure S4

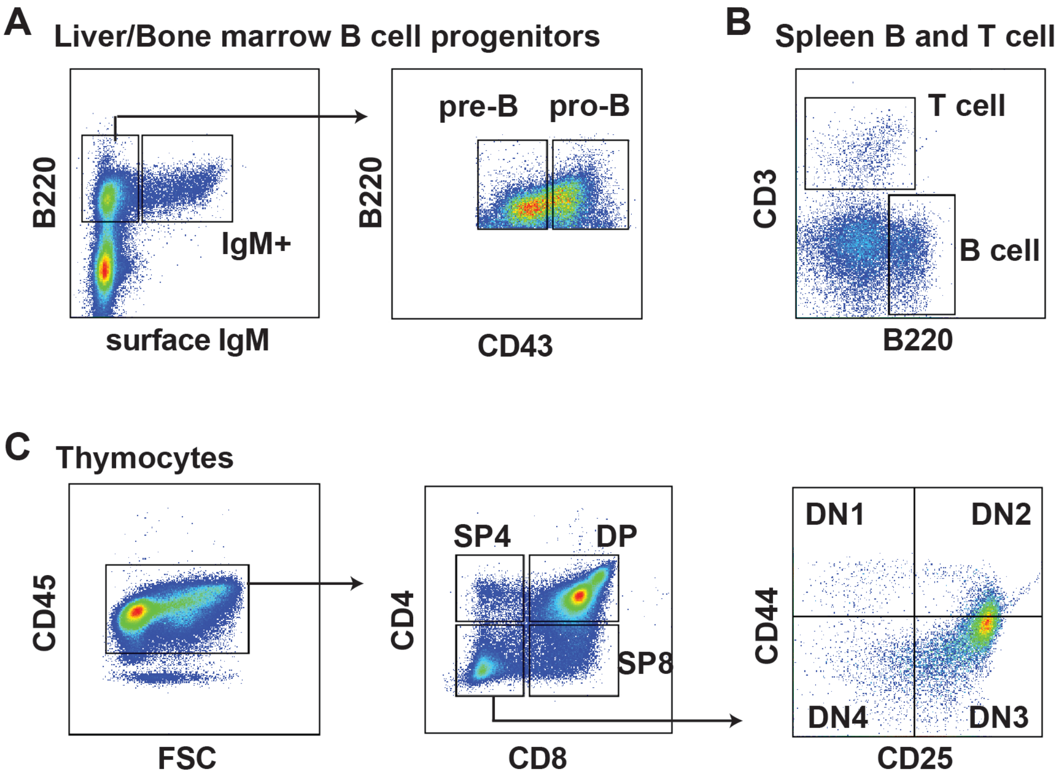


**Figure S4. Gating strategies for B cell and T cell progenitor populations.** (A) Representative gating of neonatal liver B cell progenitors. (B) Representative gating of neonatal spleen B and T cells. (C) Representative gating for neonatal thymocyte populations.

### Figure S5

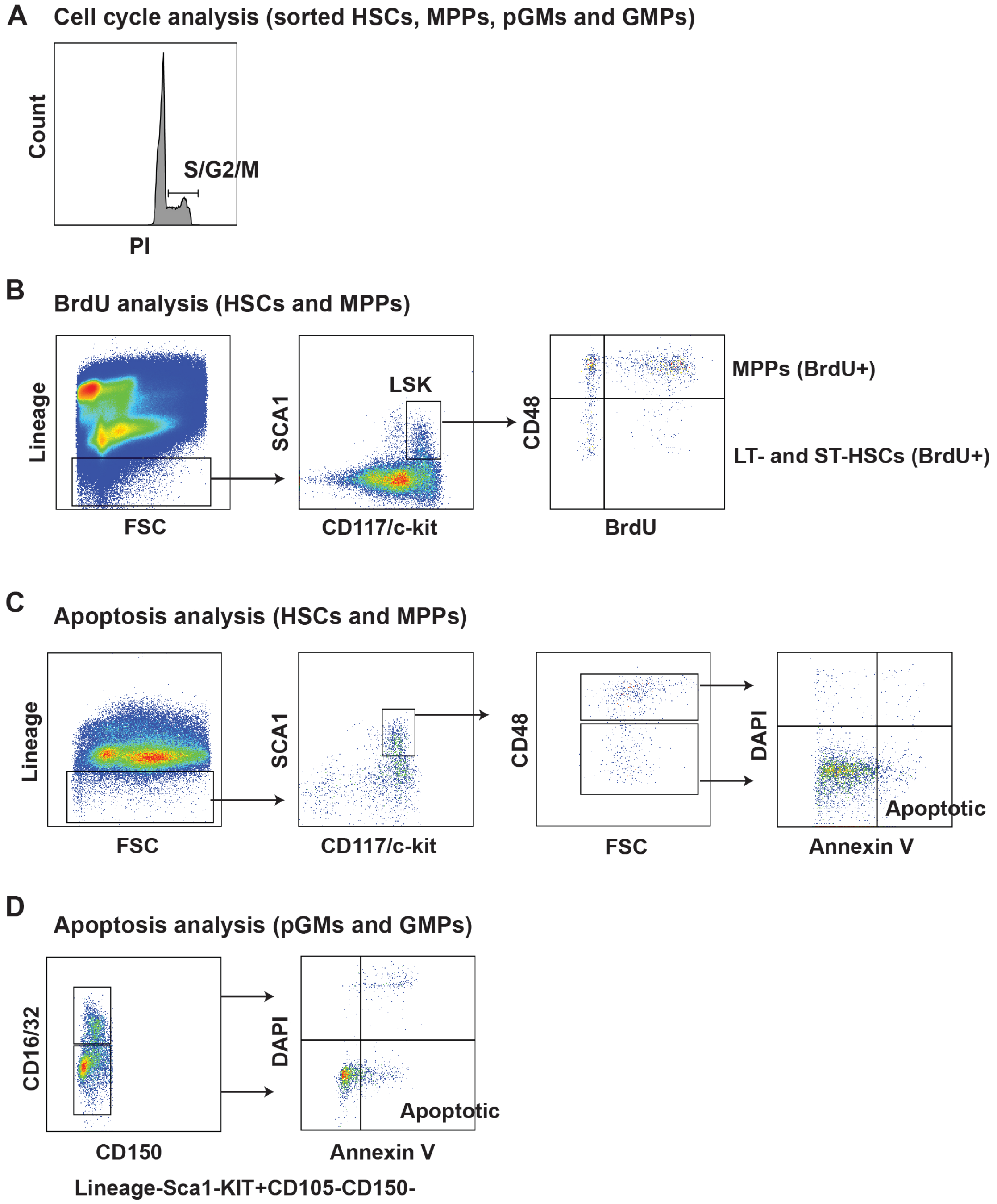


**Figure S5. Gating strategies for cell cycle, BrdU and apoptosis analyses.** (A) Representative propidium iodide stain for progenitors after sorting in 70% methanol and rehydration in PBS. (B) Representative gating strategy for neonatal BrdU incorporation assays. (C, D) Representative gating for Annexin V assays for HSCs/MPPs or myeloid progenitors. Apoptotic cells were defined as Annexin V-positive, DAPI-negative.

#### Figure S6

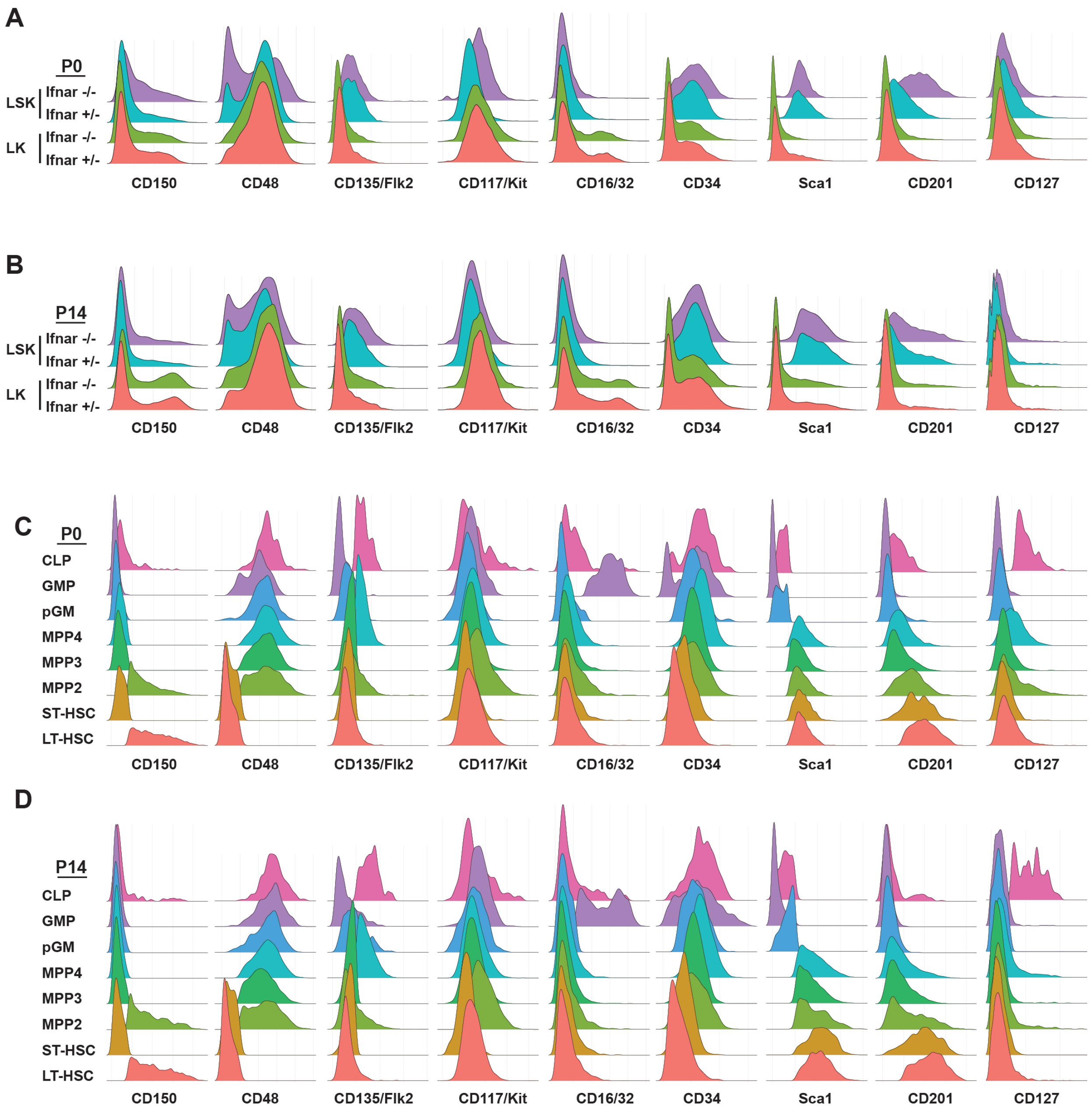


**Figure S6. Feature barcode intensities for CITE-seq experiments.** (A, B) Expression of each surface marker is shown for each sample at P0 and P14. (C, D) Expression of each surface marker is shown for each phenotypically-defined progenitor population at P0 and P14.

### Figure S7

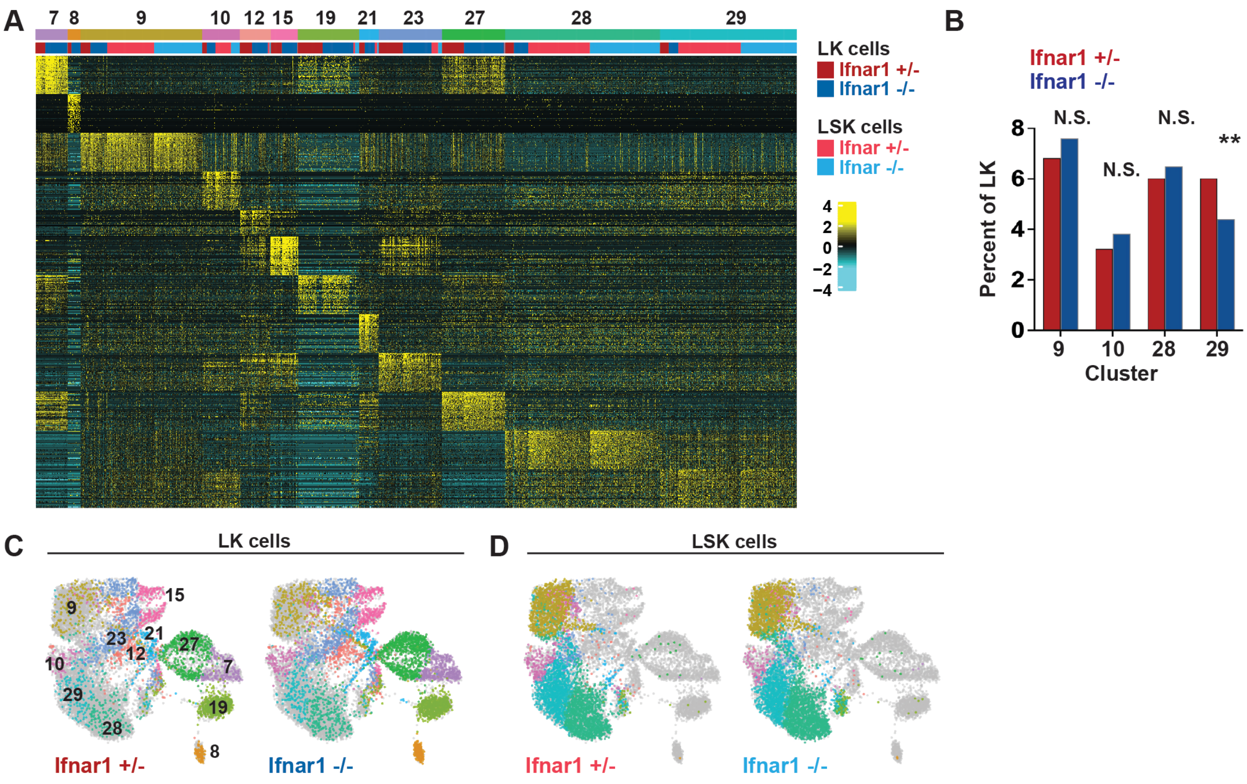


**Figure S7. CITE-seq analysis shows only very modest IFN-1-dependent changes in hematopoiesis at P14.** (A) Heatmap identifying 12 distinct clusters after iterative clustering via ICGS2. The clusters are numbered (ranging from 7-29) as defined by the ICGS2 algorithm. The clustering reflects aggregates of separately sorted LK and LSK cells from *Ifnar1+/-* and *Ifnar1-/-* mice. Genotypes and cell types are indicated by color coding in the second horizontal bar above the heatmap. (B) Percentages of LK cells in HSC/MPP clusters 9, 10, 28 and 29 in *Ifnar1+/-* and *Ifnar1-/-* mice. **p<0.01 by posthoc Chi-squared test. (C, D) UMAP plots showing clustering results for *Ifnar1+/-* and *Ifnar1-/-* LK and LSK cells. Clusters are color coded to match panel A.
